# Pangenome graph augmentation from unassembled long reads

**DOI:** 10.1101/2025.02.07.637057

**Authors:** Luca Denti, Paola Bonizzoni, Brona Brejova, Rayan Chikhi, Thomas Krannich, Tomas Vinar, Fereydoun Hormozdiari

## Abstract

Pangenomes are becoming increasingly popular data structures for genomics analyses due to their ability to compactly represent the genetic diversity within populations. Constructing a pangenome graph, however, is still a time-consuming and expensive process. A promising approach for pangenome construction consists of progressively augmenting a pangenome graph with additional high-quality assemblies. Currently, there is no method for augmenting a pangenome graph with unassembled reads from newly sequenced samples without first aligning the reads to a reference genome and performing variant calling and genotyping on the new individuals.

In this work, we present the first assembly-free and mapping-free approach for augmenting an existing pangenome graph using unassembled long reads from an individual not already present in the pangenome. Our approach consists of finding sample specific sequences in reads using efficient indexes, clustering reads corresponding to the same novel variant(s), and then building a consensus sequence to be added to the pangenome graph for each variant separately.

Using simulated reads based on Human Pangenome Reference Consortium (HPRC) assemblies, we demonstrate the effectiveness of the proposed approach for progressively augmenting the pangenome with long reads, without the need for *de novo* assembly or predicting genetic variants of the new sample. The software is freely available at https://github.com/ldenti/palss.

## 1 Introduction

The release of the first complete human reference genome sequence [29] has significantly enhanced our ability to access every region of the genome [1,30]. However, two key shortcomings of relying on a single linear reference genome have become evident: the persistent biases introduced by considering a single reference genome [28] and the inadequate representation of structural variants [20]. A single reference genome cannot capture the full genetic diversity of the human population, and studying difficult regions and complex structural variants using single linear reference genome has proven challenging. As a result, researchers are increasingly shifting towards the adoption of a *pangenome* reference, i.e., a collection of genomes that can capture the genetic variability of diverse individuals [28]. A pangenome reference is often compactly represented as a *pangenome graph*, and many approaches have been devised to take advantage of this structure and improve the accuracy of genomic analyses [36,35,9]. Improving methods for the accurate and efficient construction of pangenome graphs is an active research area. For instance, it is challenging to define the biological meaningfulness of any graph [26] and it is also not clear how an optimal pangenome graph (if it exists) should look like [3].

The most straightforward methodology to build a pangenome graph consists in embedding known variations (a VCF file) in the linear reference genome. When a new sequencing dataset is available, this requires to first map the reads and then genotype the new individual. However, the current state-of-the-art approach, `giraffe-deepvariant`;[28], does not fully address the reference bias. It relies on pangenomic information only during the read mapping step, as the graph alignments are surjected (i.e., mapped back) to the reference genome before genotyping the variations using the reference-based `;DeepVariant` [32]. Alternative approaches which tries to completely remove the reference bias start from *de novo* phased genome assemblies. Most approaches [10,11,3] require the multiple sequence alignments of the input assemblies and derive the pangenome graph from it. In contrast, an orthogonal approach [16] avoids the complexity of multiple sequence alignments but still starts from an all-vs-all alignments of the input genomes. Other approaches [27,20], instead, rely on an iterative sequence-to-graph mapping of the genomes and progressively build a pangenome graph by inserting one genome at the time. All these approaches however require *high-quality de novo* assemblies to construct pangenome graphs from sequencing data. In spite of the recent advances in long read sequencing technologies, obtaining accurate and complete *de novo* assemblies is still a costly and computationally intensive step which require high coverage data coming from different sequencing technologies [2]. On top of this complexity, most approaches also require an additional demanding preprocessing of the assemblies (multiple sequence or all-vs-all alignment) that hardly scales for many individuals. Sequence-to-graph alignment is a less demanding alternative which still, however, requires high-quality assemblies. `minigraph` [27], indeed, generalizes `minimap2` [25] minimizer-based mapping technique and can efficiently build a pangenome graph from relatively similar assemblies. However, `minigraph` includes only structural variations in the resulting graph, thus producing a lossy pangenome graph. To solve this issue, `Minigraph` has been later incorporated in the `Minigraph-Cactus` tool [20]. `Minigraph-Cactus` starts from the `minigraph` graph and augments it with Single-Nucleotide Polymorphisms (SNPs) and short indels by remapping the input assemblies to it.

Inspired by the above approaches, we propose a computational method to dynamically update (or augment) a pangenome graph starting directly from unassembled reads. Given a pangenome graph that encodes a collection of genomes and a long read sample from an individual not already represented in the collection, our goal is to update the graph topology by adding new vertices and edges to capture novel variants and potentially new sequences from the newly sequenced individual. Instead of relying on high-quality genome assemblies, we propose an assembly-free approach that is also mapping-free since it does not require to align the full reads to the input pangenome graph. Our approach targets the portions of the reads that are likely to represent novel regions of the haplotypes or new variants. After constructing the sequences corresponding to these regions, our approach incorporates them into the graph in order to fully represent the genetic diversity of the new individual. A similar idea has been recently proposed in [34], where a pangenome graph is augmented using structural variations called from graph-based long-read alignments.

Thanks to its alignment-free nature, our approach is not severely affected by mapping artifacts as alignment-based approaches. Such artifacts usually occur when aligning long reads to complex and repetitive regions [21,37]. Additionally, its assembly-free nature allows it to accurately and efficiently augment a pangenome graph even from low-coverage (10×) long-read sequencing data. Moreover, unlike `Minigraph-Cactus` that rebuilds the pangenome graph from scratch to incorporate new sequenced individuals, our method enables the updating of an existing pangenome graph with new samples without the need for a complete reconstruction. Furthermore, instead of selecting an assembly from pangenome as backbone reference and augment it with new assemblies while building the pangenome graph, our approach considers the entire existing pangenome graph as backbone and augments it accordingly with long-read sequencing data.

Our experimental evaluation, based on real pangenome graphs constructed from the Human Pangenome Reference Consortium callsets [28] and simulated HiFi reads, demonstrates the effectiveness of the proposed pangenome graph augmentation using low-coverage sequencing data from new samples. Furthermore, we show that our approach can enhance assembly-based state-of-the-art methods, such as `Minigraph-Cactus`, by expanding augmentation to regions missed during *de novo* assembly.

## 2 Preliminaries

Let *Σ* be an ordered and finite alphabet of size *σ*. In the context of DNA sequencing, we consider Σ = {*A, C, G, T, N*}. A *string* of length *n* is an ordered sequence of *n* characters drawn from *Σ*. Given a string *S*, |*S*| denotes its length and *S*[*i, j*] with 0 ≤ *i* ≤ *j <* |*S*| is the (*j* − *i* + 1)-long substring starting at position *i*. A *k*-mer is a string of fixed length *k*. When working with double-stranded DNA sequences, it is common to consider *canonical k*-mers. A canonical *k*-mer is the lexicographically smaller string between the *k*-mer itself and its reverse-and-complemented string. In the following, whenever we refer to a *k*-mer, we will implicitly refer to its canonical form.

Given two set *R* and *T* of strings, *specific strings* can be informally defined as those shortest strings occurring as substrings of some strings in *T* and not occurring in *R*. More formally, a specific string is a substring *s* of some *t* ∈ *T* that does not occur as substring of any *r* ∈ *R* while any substring of *s* occurs in some *r* ∈ *R*. For more details on this definition, its correctness, and its potential applications, we refer to [22,7]. As done in [13], we will refer to *specific superstrings* as result of merging consecutive specific strings that overlap on some *t* ∈ *T*. More precisely, we take the substring of *t* ∈ *T* starting with the first and ending with the last specific string in the list of consecutively overlapping specific strings. In the following, to simplify the discussion, we will refer to such specific superstrings as simply specific strings. Specific strings pinpoint all differences between the two input sets and can be efficiently identified using an FMD-index data structure [22] in time which is linear in the total length of the computed specific strings. The FMD-index [24] is a self-index based on the FM-index [14] which indexes a set of strings and their reverse-and-complement and allows for backward and forward extensions of a query pattern. A recent work from Li [26] improved the construction of this data structure, enabling the indexing and querying of pangenomes, encoded as a set of strings.

An alternative way to represent a pangenome is by the means of a graph structure. A *variation graph*, as defined in [6], is a directed graph *G* = (*V, E, W*) whose vertices are labeled by nonempty strings built on *Σ*, with *λ*: *V* → *Σ*^+^ being the labeling function, and where *E* denotes the set of edges and *W* denotes a nonempty set of distinguished walks (or paths). A variation graph is a compact encoding of multiple haplotypes, which are represented as distinguished paths (in this work, we will refer to these paths as *haplotype paths*). Each vertex corresponds to a portion of some haplotypes, and each edge represents consecutiveness between two such portions. We note that the haplotypes are not always readily available in all pangenome graphs constructed with current state-of-the-art approaches. In this work, we will focus on variation graphs and not sequence graphs (following the same definitions of [6]). Moreover, as an additional constraint, we will focus on acyclic variation graphs. Note that although most of our approach can also be adapted to cyclic graphs, one of its steps (i.e., the clustering of anchored specific strings) requires a topological ordering of the vertices of the graph.

## 3 Method

Given a pangenome graph and a long read sample coming from an individual not already present in the pangenome, our goal is to augment the graph in order to include the new individual. In other words, we aim at updating the graph with novel vertices and edges that represent specific portions of the haplotypes of the new individual (potentially indicating novel variants) that are not already in the pangenome. In such a way, a pangenome graph can be dynamically updated using unassembled reads, without the need for a full reconstruction.

We propose an assembly- and alignment-free method that, without requiring high-quality assembly or full sample alignments to the graph structure, targets only those portions of the reads that are specific to the individual and include them in the graph. Our approach is based on the following data-driven observation. Given a read sample sequenced from an individual not present in the collection of genomes, each read supporting any difference with the already collected genomes (e.g., sequencing errors or real variations) shows substrings describing such differences and all reads supporting the same variation contain similar substrings (similar and not equal due to sequencing errors, neighboring variations, and different ploidy). Our claim is that these substrings contain enough information to produce a local assembly of the haplotypes that can be further analyzed to augment the pangenome graph structure.

However, being able to compute and analyze these substrings in an assembly- and alignment-free fashion is not straightforward. It is first necessary to compute these substrings and cluster them by variations. Moreover, it is also necessary to place them onto the graph and then understand how to summarize the information coming from the different reads. To solve these issues, we devised an approach (outlined in Figure 1) based on *specific strings* and *graph sketching*. Specific strings pinpoint all differences with respect to the genomes encoded in the pangenome graph and the graph sketch allows to locate (*anchor* in the following) such strings onto the graph. Once all specific strings have been anchored to the graph, they are clustered based on the subgraph they have been anchored to. Each cluster represents a portion of the new individual haplotype(s) that is not already present in the graph. Each cluster is independently analyzed to produce one (or two, depending on the ploidy of the locus) consensus string(s) representing the local haplotype sequence(s) specific to the individual. These segments are finally locally aligned to the corresponding subgraph to obtain a base-level alignment that is used to augment the graph.

**Fig. 1.**
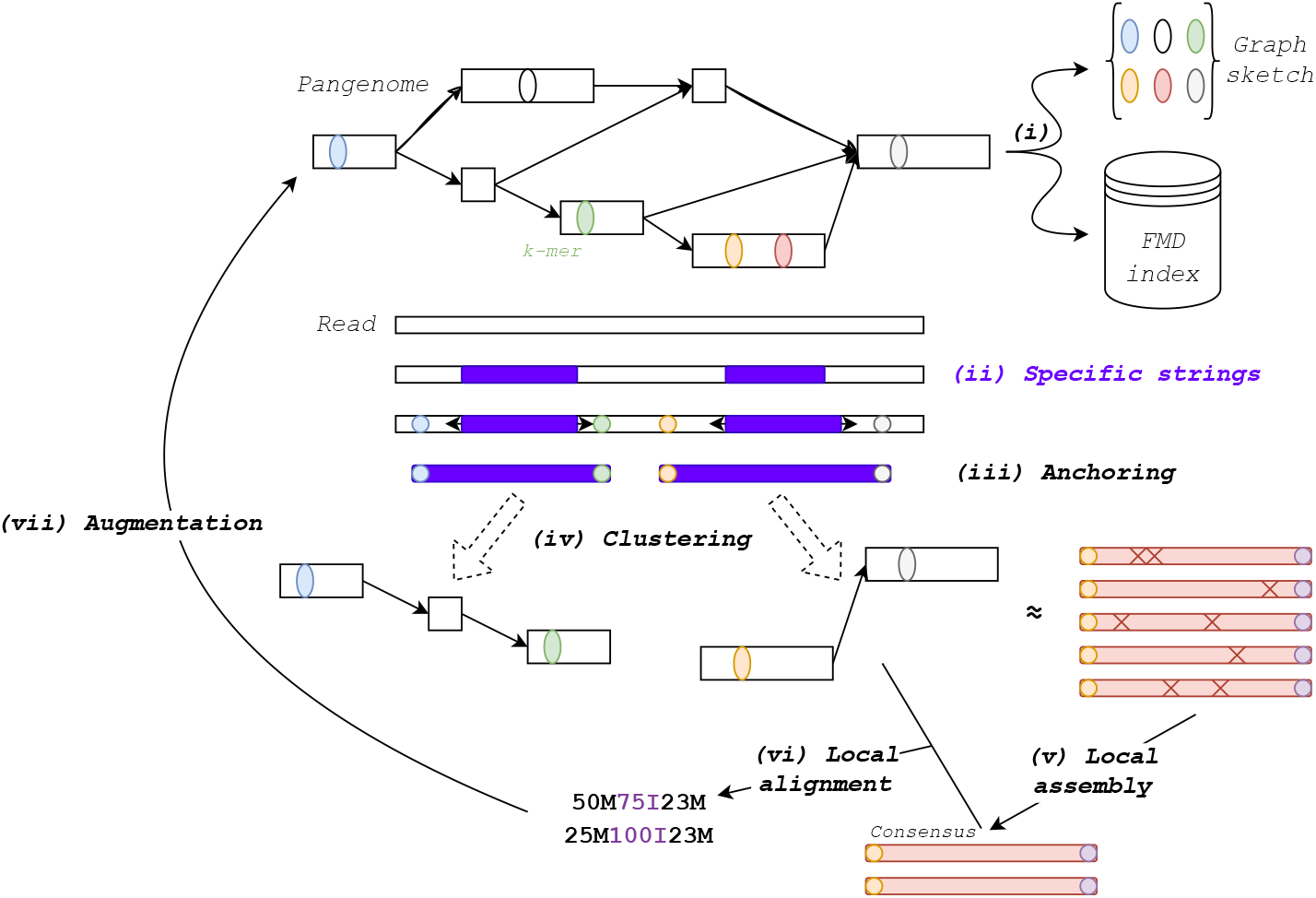
Method overview. (i) Pangenome graph is sketched using solid anchors (*k*-mers) and its distinguished paths, i.e., haplotypes paths, are indexed using an FMD-index. (ii) Specific strings are computed from long reads and (iii) anchored to the graph using solid anchors. (iv) Based on their anchoring, anchored specific strings are clustered. (v-vi) Each cluster, composed of several specific strings anchored to (mostly) the same subgraph and which may contain sequencing errors (the red Xs), is assembled and locally realigned back to the corresponding subgraph (in case of a diploid locus, two consensuses are created and analyzed). (vii) The base-level alignments are finally used to augment the initial pangenome graph.

### 3.1 Graph sketching

Inspired by other *k*-mer (and minimizer) indexes for pangenome graphs [27,5], we introduce a *k*-mer based sketch that allows to locate *k*-mers on the graph. These *k*-mers will be used to *anchor* specific strings to the graph. In more detail, anchoring a specific string to the graph means to locate the putative location in the graph (i.e., the subgraph) where the difference(s) supported by the specific string needs to be added. Let *G* = (*V, E, W*) be a variation graph representing |*W* | haplotypes. We define any *k*-mer occurring in only one vertex label (of length ≥ *k*) of *G* as an *anchor*. An anchor is unique in the vertex set, but this does not assure that the anchor is unique in the genomes. Indeed, *k*-mers overlapping the edges of the graph are not considered in this definition, i.e., any *k*-mer with a prefix matching the suffix of *λ*(*u*) and a suffix matching the prefix of *λ*(*v*), such that (*u, v*) ∈ *E*. We recall that *λ* is the labeling function, assigning a non-empty string to each vertex of the graph. To be able to retain only informative anchors, we add an additional constraint to our definition and we introduce the notion of *solid anchor*. A solid anchor is any anchor occurring ≤ |*W* | times in the collection of haplotypes. This heuristically-derived definition allows us to keep all those anchors that can be used to uniquely anchor a specific string on the graph while filtering out anchors with too many occurrences in the genomes. In short, a *graph sketch* will consist of the set of all solid anchors of the graph, where each anchor *a* is stored alongside a pair (*a*_*v*_, *a*_*o*_) indicating the vertex *v* associated to *a* and the offset *o* from the starting position of *λ*(*v*).

### 3.2 Specific strings and their anchoring

To find specific strings representing the differences between the read sample and the pangenome, we index the set of genomes encoded in the pangenome graph with an FMD-index. Using this index, specific strings are computed using the algorithm described in [22], which we reimplemented to work with the new `ropebwt3` implementation of FMD-index [26]. We use error-corrected reads to reduce the number of specific strings, leading to improved precision and reduced running time. Specific strings need to be placed onto the graph, which is not trivial, as they are defined as strings that are not present in the pangenome index. To overcome this problem, we rely on the graph sketch and solid anchors described in the previous section. In particular, we search the graph sketch for *k*-mers upstream and downstream the specific string on the read. More formally, let *x* = *R*[*b, e*] be a specific string contained in read *R* of length *r*. We check the upstream *k*-mers *R*[*i, i* + *k*] with 0 ≤ *i* ≤ *b* − *k* and downstream *k*-mers *R*[*j, j* + *k*] with *e* ≤ *j* ≤ *r* − *k*. If a *k*-mer is found as a solid anchor in the graph sketch, we retrieve the corresponding vertex and offset. Ideally, we would select a pair of upstream and downstream solid anchors that are located in the correct order on a haplotype path of the pangenome graph and are as close as possible to each other. However, this would require searching through paths of the graph, which is computationally intensive. To simplify matters, we instead use a simple heuristic. We number the vertices of the pangenome in the topological ordering and we select the pair of anchors upstream and downstream the specific string with the smallest difference in vertex number and in the correct order. In case of a tie, i.e., pairs of anchors coming from the same two vertices, we select the pair closest to the specific string on the read sequence. To further reduce the complexity of this step, we limit our computation to 20 solid anchors upstream and 20 solid anchors downstream (the number is a user-defined parameter). Moreover, to reduce the likelihood of misplacing an anchor due to a novel repeat in the read sequence (for instance in presence of a duplication), we limit our search to those *k*-mers that are unique in the read.

The result of this computation is a substring of a read that starts and ends with solid anchors, whose locations in the graph are known. We call such a substring an *anchored specific string*.

### 3.3 Clustering of anchored specific strings

The anchored specific strings need to be sorted into clusters which represent the same genomic locus. Here, the genomic locus of an anchored specific string can be represented by the subgraph composed of all paths connecting the two anchors. The clustering then could join all specific strings whose subgraphs overlap.

To speed up the clustering, we approximate each subgraph as an interval over the topological ordering of the vertices. More formally, given a specific string anchored at vertices with order *a* and *b* (with *a* ≤ *b*) in the topological ordering, we represent the subgraph corresponding to the string simply by interval ⟨*v*_*a*_, *v*_*b*_. ⟩Two anchored specific strings are then clustered together if their intervals overlap. This representation allows us to find clusters by a simple and efficient sweep-line algorithm. We first sort the specific strings based on the vertex and offset of their upstream anchor, i.e., we sort them by their starting position along the pangenome graph. The first specific string forms a cluster, and all subsequent specific strings are compared to the last created (or updated) cluster. In more detail, we iterate over the sorted list and we progressively add the current specific string to the last created cluster if intervals overlap or we create a new cluster containing only the last specific string. At each iteration, we keep only one open cluster and, when we add a new specific strings to it, we also maintain the left-most and right-most anchors.

Note that this heuristic representation may sometimes cluster together strings whose true subgraphs do not actually overlap since an interval of the topological ordering may contain vertices not reachable from the two anchors (e.g., in the presence of superbubbles). However, in practice, this does not affect the accuracy of our heuristic for clustering specific strings. Indeed, we aim at grouping specific strings that are relatively close: each cluster will be successively refined and the additional vertices will not be considered while augmenting the graph.

### 3.4 Cluster analysis

At this point, each cluster represents a locus of the new genome exhibiting some sort of difference with the set of genomes encoded in the pangenome. We need now a ploidy-aware way to summarize the information coming from the different reads. Indeed, since we clustered the anchored specific strings only based on their position along the pangenome graph, a cluster may contain specific strings coming from multiple haplotypes, e.g., in case of multi-allelic variations.

We start by filtering out those clusters likely generated by sequencing errors, i.e., we consider only those clusters supported by specific strings coming from at least *s* = 2 (user-defined parameter) reads. We recall that a cluster is a set of anchored specific strings where each specific string starts and ends with solid anchors. However, due to the clustering step, different anchored specific strings in the same cluster may start and end with different anchors. To simplify the analysis of each cluster, we further extend the specific strings to make them start and end with the same solid anchors. While performing the clustering, we kept track of the left-most and right-most solid anchors of each cluster and, therefore, we can now extend each anchored specific string up to these anchors, when possible. If not possible, the specific string is filtered out (we note that, in practice, this does not affect the accuracy of our augmentation). In such a way, we force each cluster to be composed of anchored specific strings that start and end with the same solid anchors. We are now ready to analyze all the resulting clusters supported by at least *s* reads, one by one.

We first split a cluster based on the length of its specific strings to handle multi-allelic variations of different lengths. Every time the ratio between the length of two anchored specific strings is above a threshold (0.97 by default), they are considered as coming from the same haplotype and then clustered together. Indeed, in the presence of multi-allelic indels or structural variations, we should expect specific strings of different length. However, this additional (sub)clustering step does not work for SNPs or other variations with alleles of similar length. We will handle these in the next step.

Once the cluster is split by specific string length, we consider the *h* most supported (sub)clusters (*h* = 2 in case of diploid individual) and we compute a consensus from each of them via partial order alignment of their specific strings. To do so, we rely on the `abpoa` library [15]. If we already have two subclusters (created based on the length of the specific strings), we build a single consensus for each of them using `abpoa`. If we have a single subcluster, instead, we create a maximum of two consensus, depending on their similarity (we may indeed be in a heterozygous locus or a multi-allelic locus whose variations cannot be simply discerned by their size only). To do so, we ran `abpoa` enabling the generation of multiple consensus (by setting `max_n_cons` to 2 and `min_freq` to 0.25, when needed). We note that in the case of homozygous events, we expect the partial order alignment to produce a single consensus.

All consensus created from at least *s* specific strings are then aligned back to the local subgraph induced by the cluster since they potentially represent novel sequences that need to be added to the graph. Since we do not want to model recombination among the input genomes, we decided to align via global dynamic programming the consensus to the haplotype paths of the subgraph and then keep the alignment with the best alignment score. Since, as described above, the subgraphs may contain vertices not reachable from the two solid anchors characterizing the considered cluster, we consider only those haplotype paths going through both their vertices. Moreover, since multiple haplotype paths may result in the same genomic sequence, we first merge all paths based on the sequence they induce and then we align the consensus to this subset of paths. To perform the global alignment, we use the `ksw2` library [25]. This is the main bottleneck of the approach since it requires to align the same consensus to multiple paths. Future works will be devoted to extend our approach to align the consensus directly to the subgraph via dynamic programming, in a similar way as proposed in [4].

Finally, the information provided by the base alignment of each consensus is used to augment the graph. By analyzing the difference string computed during the alignment to a path of the graph, it is possible to embed all differences into the graph by splitting original vertices and adding new vertices and edges. To perform this step, we rely on `vg augment` [19] which embeds variations from graph alignments into the graph.

## 4 Results

To evaluate the accuracy of our method, we considered pangenome graphs constructed from the variant callsets against the T2T reference genome provided by the Human Pangenome Reference Consortium (HPRC) [28] using `vg` [17]. For various experiments, we used different numbers of randomly chosen individuals *n* ∈ {1, 2, 4, 8, 16, 32}. Note that a pangenome for *n* individuals contains 2*n* + 1 haplotypes (two haplotypes per individual plus the T2T reference genome). Duplicated variations were removed from the callset, following the definition used in the `vg` toolkit, i.e., two variants are duplicated if their chromosome, position, and alleles are the same. As required by individual experiments, these pangenomes were further augmented with our method using simulated reads from a randomly chosen new individual not included in the pangenome. Code and instructions to reproduce our experimental evaluation are available at https://github.com/ldenti/palss. Running times and memory requirements of our approach are reported in Supplementary Section A.

### 4.1 Most of the genome is well covered by solid anchors

Our method uses only reads that contain at least two solid anchors. These anchors are used to place specific strings in the graph and to cluster them. In our first experiment, we verified that our solid anchors are sufficient to anchor specific strings in the vast majority of the human genome using long-read data. We built a sketch of the solid anchors (with *k* = 27) for a pangenome graph consisting of three chromosomes (1, 11, and 20) from 32 individuals. We then searched for regions in the T2T reference where the solid anchors are more than 15kbp apart, which is a typical average read length for PacBio HiFi reads. These regions are thus inaccessible to our method.

There are 488 of such regions which account for 3.4% of the three chromosomes considered (4.8% of chromosome 1, 1.0% of chromosome 11, and 2.6% of chromosome 20). Size distribution of these regions is reported in Figure 2a. All of these regions intersect known challenging regions of the genome, namely, simple repeats, satellites, LINEs, centromeres, and segmental duplications. The longest one, almost 200kbp in length, is located inside the centromere of chromosome 1 (chr1: 125983206-126180483).

**Fig. 2.**
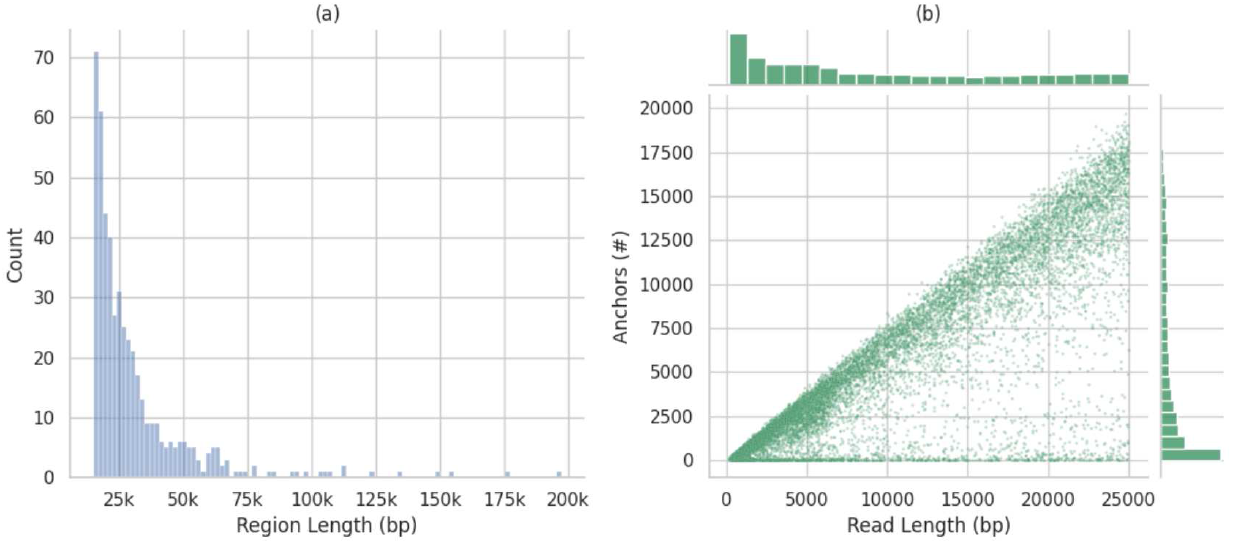
Coverage of the genome by solid anchors. (a) Size distribution of the 488 regions longer than 15kbp without any solid anchor. (b) Number of anchors and read length distribution in Oxford Nanopore reads. We restricted this plot to all reads shorter than 25kbp. The full version of this plot is reported in Supplementary Figure A.

We also evaluated if our solid anchors can be actually found in real sequencing reads. We considered a set of Rl0.4 simplex Oxford Nanopore sequencing reads amounting to 10 x coverage of chromosome 20 [23]. Out of 26 198 reads, 99.4% had at least two anchors, 99% had at least seven anchors, and 98% at least 26 anchors; in fact the number of anchors is well correlated with the read length (Figure 2b). We note that we perform this analysis without correcting the ONT reads.

Overall, this analysis shows that except for a few regions that are notoriously difficult to analyze, our solid anchors are sufficient to anchor sample-specific strings in the vast majority of the human genome using long-read data.

### 4.2 Graph augmentation improves read mapping

In this section we demonstrate that our graph augmentation yields graphs that capture novel variants present in the added individual. To this end, we first constructed a pangenome graph of chromosome 20 from n individuals (with n E {1, 2, 4, 8, 16, 32}). We used an additional individual to augment this graph with our method using simulated HiFi sequencing reads, which are obtained from the two haplotypes of the new individual using `pbsim3` **[31]** with 5 x coverage per haplotype. The reads are error-corrected by `hifiasm` [8] prior to running our algorithm (using solid anchors of size k = 27). For comparison purposes, we also created a graph directly from n *+* 1 individuals (using vg), including the new individual used for augmentation. This graph is denoted as `FULL`, the graph built from n individuals is denoted as `10UT,` and the augmented graph as `10UT-AUG.` To put our results in perspective, we also created a pangenome graph with Minigraph-Cactus (graph `MGC`). We defer the discussion of this comparison to Section 4.4.

Figure 3 demonstrates that the simulated reads can be aligned to the augmented graph much better than to the `10UT` graph. In particular, we aligned the simulated reads using minigraph [27].

**Fig. 3.**
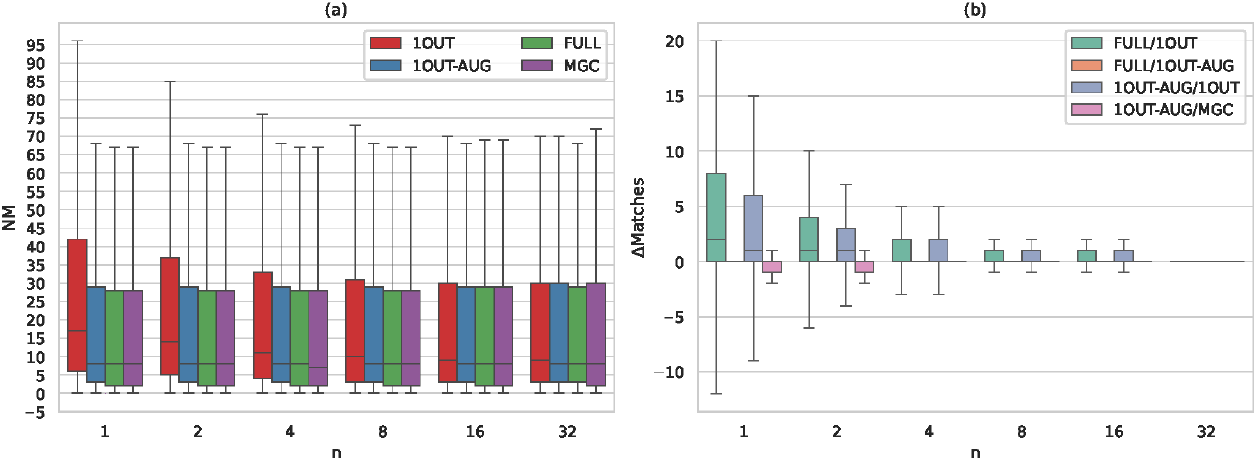
Results on alignment accuracy (alignments covering more than 80% of the reads). (a) Distri bution of the edit distances reported by `minigraph` (NM field) when aligning simulated reads to the 4 graphs considered in our analysis (lower is better). (b) Pairwise comparison of the alignments in terms of difference in the number of residue matches *(ΔMatches)* reported by `minigraph.` In each plot, the results are broken down by the number n of individuals included in the original pangenome graph (x axis). For the sake of clarity, outliers are not plotted. `FULL`: graph built from n + l individuals, `1OUT`: graph built from n individuals, `1OUT-AUG: 1DUT` augmented with the new individual, MGC: assembly-based graph built by `; Minigraph-Cactus` (Section 4.4).

Note that `vg` can create cycles during augmentation, and thus we are limited to long read aligners that can accommodate cyclic graphs. We only considered alignments covering more than 80% of each read and, in case of multiple alignments for the same read, we selected the one with the highest ratio between the number of residue matches and alignment length. For pangenomes constructed from a small number of individuals, there is a large difference between median edit distance of the reads aligned to the 10UT graph compared to the full graph, caused by variations specific to the new individual (Figure 3(a)). The edit distance of the reads aligned to the aug mented graph is very close to that of the full graph, suggesting that these variants are for the most part captured by our method. As the size of the pangenome increases, the differences be tween graphs become smaller, which is expected, since the number of novel variants contributed by the new individual diminishes.

A more detailed comparison between pairs of graphs is shown in Figure 3(b) which depicts the distribution of the difference between the number of residue matches for individual reads aligned to two different graphs. The median difference between the number of matches to `FULL` and `10UT-AUG` is zero in all cases. As before, the difference between `; FULL` and `10UT` decreases with the number of individuals included. Surprisingly, some reads exhibit a higher number of matches when aligned to the `10UT` graph instead of the more complete `FULL` and `10UT-AUG.` We suspect that the presence of complex subgraphs consisting of several close bubbles may affect `minigraph` alignments accuracy, forcing it to produce partial alignments. Indeed, `minigraph` is primarily designed to work with graphs that include mostly structural variations. When we consider only reads fully aligned, the number of reads aligned better to the `10UT` graph drops significantly (Supplementary Figure B). We also ran the same analysis using GraphAligner [33] instead of `minigraph` (results described in Supplementary Section B) but we did not observe any significant change in terms of alignments accuracy (although `GraphAligner` produces less clipped alignments).

### 4.3 Graph augmentation is precise

To evaluate the precision of our augmentation, we analyzed if the new vertices added to the input graph are actually used by the alignments. Although not perfect, since some reads may not be fully aligned by `minigraph,` this analysis provides an upper bound to the number of false additions our augmentation made. As expected, the lower the number of individuals included in the graph, the higher the number of new vertices added to the graph during its augmentation (Figure 4a). Remarkably, almost all the novel vertices (9 9% whenn = 1 and more than 95% with n = 4) are covered by at least one alignment.

**Fig. 4.**
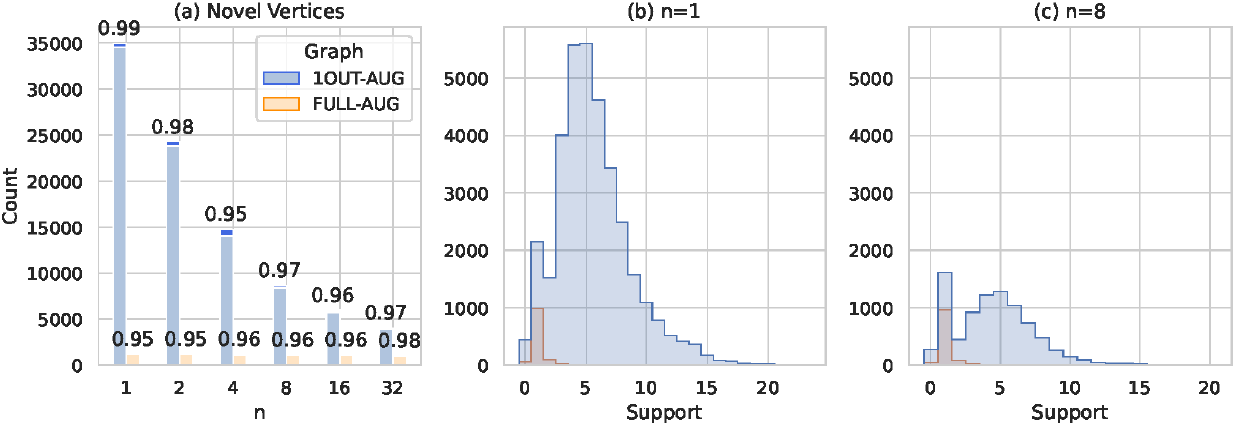
Results on augmentation precision. (a) The number of novel vertices added by our augmentation from long reads in the `1OUT-AUG` and `; FULL-AUG` graphs are shown. Lighter bars represent the total number of vertices supported by at least one alignment whereas darker bars represent the number of vertices not covered by any alignment. The number on top of each bar is the percentage of vertices supported by at least one alignment over the total number of novel vertices. (b-c) Distribution of the support (number of alignments) of each novel vertex added to pangenome graph by the augmentation method to the `1OUT-AUG` and `FULL-AUG` graphs (same color coded as (a))

As an additional control, we also used our augmentation on the `FULL` graph, which already includes all variation from the new individual and thus ideally, no new vertices would be added, resulting in a graph denoted `FULL-AUG.` Indeed, the number of new vertices added to `FULL-AUG` is much lower than in the case of `10UT-AUG,` and surprisingly, these new vertices are also mostly supported by the alignments. However, the vast majority of these vertices are covered by just one read alignment (Figure 4(b,c)), suggesting that they could represent sequencing errors wrongly mistaken for real variations.

A similar trend can be seen when considering the vertices with support one added to the `10UT` graph. In this case, there are more of these vertices added to the n = 1 graph instead of then = 8 graph (2144 out of 34 968 versus 1610 out of 8640). Although we suspect that some of these vertices are actually real variations that are not covered by `minigraph` alignment due to clipped alignments, most of these are actually sequencing errors, wrongly included in the new graph (the same trend can be observed when considering the alignments computed with `GraphAligner` Supplementary Section B). Future works will focus in improving our method to better discern sequencing errors left in the reads from real variations. Note that, the median and peak support for novel vertices added to the 10UT graph is at half the sequencing coverage (i.e., 5 x) providing additional evidence of the precision of our approach and the fact that most of the novel vertices added to graph are heterozygous variations.

### 4.1 Comparison to Minigraph-Cactus

To put our results in perspective, we also considered a more traditional approach of incorporating a new individual, namely assembling the new individual’s genome first using `hifiasm` and then building the pangenome of all n *+* 1 individuals using `Minigraph-Cactus.` This graph is denoted as `MGC.` Supplementary Table B reports the quality assessment of the assembly computed with quast [18] (then known genomes are obtained directly from the HPRC callset using bcftools consensus [12]). When comparing `10UT-AUG` and `MGC` graphs, read alignments to `MGC` graph are slightly better on average (Figure 3(b)), although few reads are aligned better to the `10UT-AUG` graph. However, we believe that these results may be marginally biased towards the `MGC` graphs, since they are built by aligning input assemblies with `minigraph` itself.

The main advantage of our approach is that it does not require high-quality assemblies to correctly augment a pangenome graph, and thus it should work even in those regions where the assembler has not been able to produce any contig. To verify this, we manually investigated the 35 regions (accounting for 0.16% of the chromosome) not covered by the assembly produced by `hifiasm` for the new individual. We discovered 15 regions with 65 variations specific to the new individual compared to the `10UT` pangenome for *n* = 1. Some of these regions are challenging telomeric and centromeric regions while most of them are covered by fewer than 2 reads per haplotype (aligned to the reference with `minimap2` [25]). In the remaining regions, our approach has been able to correctly augment the graph with 10 variations (SNPs and short indels) that have been missed by `Minigraph-Cactus.`

As an example, we consider the `chr20: 17563000-17563300` locus (Figure 5). In this region, as per HPRC VCF, there are a lbp insertion (A) and a SNP (C>T) that are specific to the new individual (`;HG01123,` both variations are homozygous alternative, as shown in Figure 5(a)) and are not present in the individual included in the pangenome (`;HG01361`). This is confirmed by the initial graph (`;10UT,` Figure 5(b)) which does not contain any information about these variations. The same can be seen in the `MGC` graph (Figure 5(c)): a single long vertex encompassing this genomic locus. On the other hand, after our augmentation from long reads, the new graph (`;10UT-AUG,` Figure 5(d)) contains an additional bubble (representing the C>T SNP that is specific to the new individual) and a new lbp-long vertex (representing the insertion). This is a clear example where an approach requiring high-quality assemblies is not able to build the correct graph from low-coverage sample while our approach using raw reads and specific strings is able to correctly augment the graph.

**Fig. 5.**
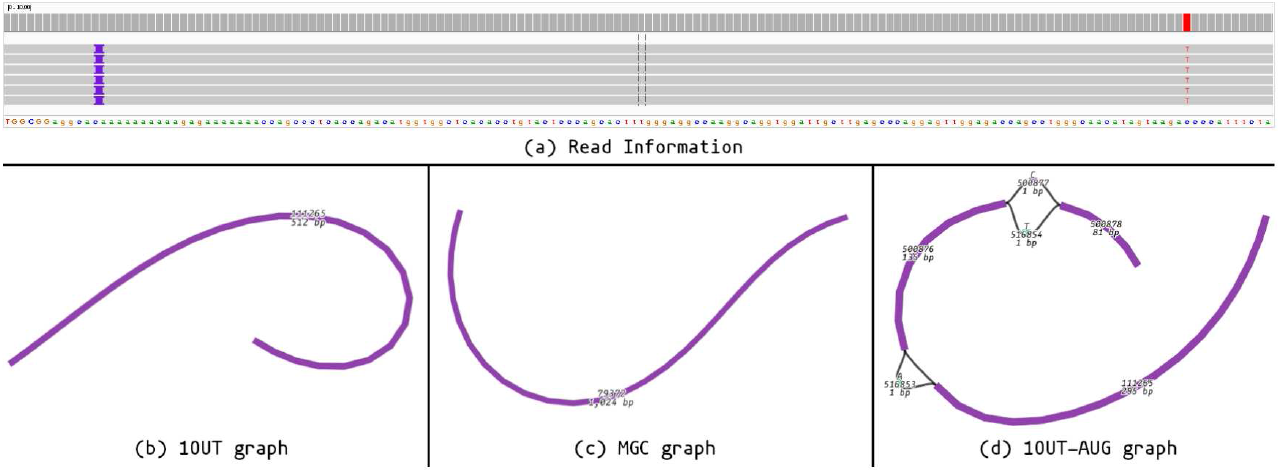
Example of augmentation. (a) Reads aligned to T2T reference genome. (b) Original graph (`1OUT`;) does not contain the two variations supported by the reads. (c) Graph built with Minigraph-Cactus also does not contain the two variations. (d) Augmented graph (`;1OUT-AUG`) includes a new bubble (the C>T SNP) and a new lbp-long vertex labeled A (the insertion) that break the initial vertex 111265 (of `1OUT` graph). Each vertex is labeled with its unique identifier and the length of its sequence. The bubbles in (d) are also labeled with the vertex sequences.

## 5 Conclusions

In this work, we addressed the challenge of dynamically updating a pangenome graph with new sequences (and variants) from a recently high-quality sequenced sample. Unlike the ex isting state-of the-art methods, we propose a novel assembly- and mapping-free approach that enables direct augmentation of the pangenome graph from long-read data, eliminating the need for accurate *de nova* assembly. Our results with simulated data demonstrate that our augmen tation approach is both accurate and comparable to state-of the-art assembly-based methods for augmenting pangenome graphs. Furthermore, we have shown that our method can successfully augment a pangenome graph even in regions missed by state-of the-art assembly based approach, `Minigraph-Cactus.` Anyway, we point out one more time that our approach requires high-quality (or error corrected) reads. Future works will be devoted to relax this constraint by improving the overall accuracy of our augmentation or by providing additional strategies to polish low-coverage long-reads samples. In addition to augmenting a pangenome graph using unassembled reads, our method can be extended to other applications. Specifically, our approach could be used to fill in missing sequences in samples already included in the pangenome graph. This would enhance the accuracy and completeness of the genomic representations in the pangenome.

We acknowledge that the challenge of augmenting a pangenome graph directly from read data is not yet fully resolved. Our primary goal is to enhance the scalability of the proposed method and enable it to work with pangenome graphs that contain cycles. Furthermore, this work presents the essential method for the first major progress in the dynamic construction of pangenome graphs. Our approach augment the graph with new vertices and edges but it does not yet update the paths within the graph. As a result, while the new individual is represented in the graph, its haplotypes (or portions thereof) are not actually included as new paths in the graph. Finally, we plan to develop an extension of the proposed method that enables the simultaneous detection of novel variants, including structural variants, in comparison to pangenome references, while augmenting the pangenome graph.

## Acknowledgments

This research work has received funding from the European Union’s Horizon programme under the Horizon Europe grant agreement (ASVA-CGR No. 101180581 to L.D.). This work has also been supported in part by National Science Foundation (NSF) award DBI-2042518 to F.H. This research was supported in part by the Slovak Grant Agency VEGA grants 1/0538/22 (T.V.) and 1/0140/25 (B.B.). P.B. has received funding from the European Union’s Horizon 2020 research and innovation programme under the Marie Skłodowska-Curie grant agreement PANGAIA No. 872539 and ITN ALPACA N.956229 and from Next Generation EU - Mission 4, MIUR 2022YRB97K, PINC, Pangenome Informatics: from Theory to Applications.

## References

1. Aganezov, S., Yan, S.M., Soto, D.C., Kirsche, M., Zarate, S., Avdeyev, P., Taylor, D.J., Shafin, K., Shumate, A., Xiao, C., et al.: A complete reference genome improves analysis of human genetic variation. Science 376(6588), eabl3533 (2022)

2. Antipov, D., Rautiainen, M., Nurk, S., Walenz, B.P., Solar, S.J., Phillippy, A.M., Koren, S.: Verkko2: Integrating proximity ligation data with long-read de bruijn graphs for efficient telomere-to-telomere genome assembly, phasing, and scaffolding. bioRxiv pp. 2024–12 (2024)

3. Avila Cartes, J., Bonizzoni, P., Ciccolella, S., Della Vedova, G., Denti, L.: Pangeblocks: customized construction of pangenome graphs via maximal blocks. BMC bioinformatics 25(1), 344 (2024)

4. Avila Cartes, J., Bonizzoni, P., Ciccolella, S., Della Vedova, G., Denti, L., Didelot, X., Monti, D.C., Pirola, Y.: Recgraph: recombination-aware alignment of sequences to variation graphs. Bioinformatics 40(5), btae292 (2024)

5. Aylward, A.J., Petrus, S., Mamerto, A., Hartwick, N.T., Michael, T.P.: Pankmer: k-mer-based and reference-free pangenome analysis. Bioinformatics 39(10), btad621 (2023)

6. Baaijens, J.A., Bonizzoni, P., Boucher, C., Della Vedova, G., Pirola, Y., Rizzi, R., Sirén, J.: Computational graph pangenomics: a tutorial on data structures and their applications. Natural Computing 21(1), 81–108 (2022)

7. Béal, M.P., Crochemore, M.: Fast detection of specific fragments against a set of sequences. In: International Conference on Developments in Language Theory. pp. 51–60. Springer (2023)

8. Cheng, H., Concepcion, G.T., Feng, X., Zhang, H., Li, H.: Haplotype-resolved de novo assembly using phased assembly graphs with hifiasm. Nature methods 18(2), 170–175 (2021)

9. Ciccolella, S., Cozzi, D., Della Vedova, G., Kuria, S.N., Bonizzoni, P., Denti, L.: Differential quantification of alternative splicing events on spliced pangenome graphs. PLOS Computational Biology 20(12), e1012665 (2024)

10. Colquhoun, R.M., Hall, M.B., Lima, L., Roberts, L.W., Malone, K.M., Hunt, M., Letcher, B., Hawkey, J., George, S., Pankhurst, L., et al.: Pandora: nucleotide-resolution bacterial pan-genomics with reference graphs. Genome biology 22, 1–30 (2021)

11. Dabbaghie, F., Srikakulam, S.K., Marschall, T., Kalinina, O.V.: Panpa: generation and alignment of panproteome graphs. Bioinformatics Advances 3(1), vbad167 (2023)

12. Danecek, P., Bonfield, J.K., Liddle, J., Marshall, J., Ohan, V., Pollard, M.O., Whitwham, A., Keane, T., McCarthy, S.A., Davies, R.M., et al.: Twelve years of samtools and bcftools. Gigascience 10(2), giab008 (2021)

13. Denti, L., Khorsand, P., Bonizzoni, P., Hormozdiari, F., Chikhi, R.: Svdss: structural variation discovery in hard-to-call genomic regions using sample-specific strings from accurate long reads. Nature Methods 20(4), 550–558 (2023)

14. Ferragina, P., Manzini, G.: Opportunistic data structures with applications. In: Proceedings 41st annual symposium on foundations of computer science. pp. 390–398. IEEE (2000)

15. Gao, Y., Liu, Y., Ma, Y., Liu, B., Wang, Y., Xing, Y.: abpoa: an simd-based c library for fast partial order alignment using adaptive band. Bioinformatics 37(15), 2209–2211 (2021)

16. Garrison, E., Guarracino, A., Heumos, S., Villani, F., Bao, Z., Tattini, L., Hagmann, J., Vorbrugg, S., Marco-Sola, S., Kubica, C., et al.: Building pangenome graphs. Nature Methods pp. 1–5 (2024)

17. Garrison, E., Sirén, J., Novak, A.M., Hickey, G., Eizenga, J.M., Dawson, E.T., Jones, W., Garg, S., Markello, C., Lin, M.F., et al.: Variation graph toolkit improves read mapping by representing genetic variation in the reference. Nature biotechnology 36(9), 875–879 (2018)

18. Gurevich, A., Saveliev, V., Vyahhi, N., Tesler, G.: Quast: quality assessment tool for genome assemblies. Bioinformatics 29(8), 1072–1075 (2013)

19. Hickey, G., Heller, D., Monlong, J., Sibbesen, J.A., Sirén, J., Eizenga, J., Dawson, E.T., Garrison, E., Novak, A.M., Paten, B.: Genotyping structural variants in pangenome graphs using the vg toolkit. Genome biology 21, 1–17 (2020)

20. Hickey, G., Monlong, J., Ebler, J., Novak, A.M., Eizenga, J.M., Gao, Y., Marschall, T., Li, H., Paten, B.: Pangenome graph construction from genome alignments with minigraph-cactus. Nature biotechnology 42(4), 663–673 (2024)

21. Jain, C., Rhie, A., Hansen, N.F., Koren, S., Phillippy, A.M.: Long-read mapping to repetitive reference sequences using winnowmap2. Nature Methods 19(6), 705–710 (2022)

22. Khorsand, P., Denti, L., Consortium, H.G.S.V., Bonizzoni, P., Chikhi, R., Hormozdiari, F.: Comparative genome analysis using sample-specific string detection in accurate long reads. Bioinformatics advances 1(1), vbab005 (2021)

23. Kolesnikov, A., Cook, D., Nattestad, M., Brambrink, L., McNulty, B., Gorzynski, J., Goenka, S., Ashley, E.A., Jain, M., Miga, K.H., et al.: Local read haplotagging enables accurate long-read small variant calling. Nature Communications 15(1), 5907 (2024)

24. Li, H.: Exploring single-sample snp and indel calling with whole-genome de novo assembly. Bioinformatics 28(14), 1838–1844 (2012)

25. Li, H.: Minimap2: pairwise alignment for nucleotide sequences. Bioinformatics 34(18), 3094–3100 (2018)

26. Li, H.: Bwt construction and search at the terabase scale. arXiv preprint arXiv:2409.00613 (2024)

27. Li, H., Feng, X., Chu, C.: The design and construction of reference pangenome graphs with minigraph. Genome biology 21, 1–19 (2020)

28. Liao, W.W., Asri, M., Ebler, J., Doerr, D., Haukness, M., Hickey, G., Lu, S., Lucas, J.K., Monlong, J., Abel, H.J., et al.: A draft human pangenome reference. Nature 617(7960), 312–324 (2023)

29. Nurk, S., Koren, S., Rhie, A., Rautiainen, M., Bzikadze, A.V., Mikheenko, A., Vollger, M.R., Altemose, N., Uralsky, L., Gershman, A., et al.: The complete sequence of a human genome. Science 376(6588), 44–53 (2022)

30. Olson, N.D., Wagner, J., Dwarshuis, N., Miga, K.H., Sedlazeck, F.J., Salit, M., Zook, J.M.: Variant calling and benchmarking in an era of complete human genome sequences. Nature Reviews Genetics 24(7), 464–483 (2023)

31. Ono, Y., Hamada, M., Asai, K.: Pbsim3: a simulator for all types of pacbio and ont long reads. NAR Genomics and Bioinformatics 4(4), lqac092 (2022)

32. Poplin, R., Chang, P.C., Alexander, D., Schwartz, S., Colthurst, T., Ku, A., Newburger, D., Dijamco, J., Nguyen, N., Afshar, P.T., et al.: A universal snp and small-indel variant caller using deep neural networks. Nature biotechnology 36(10), 983–987 (2018)

33. Rautiainen, M., Marschall, T.: Graphaligner: rapid and versatile sequence-to-graph alignment. Genome biology 21(1), 253 (2020)

34. Schloissnig, S., Pani, S., Rodriguez-Martin, B., Ebler, J., Hain, C., Tsapalou, V., Söylev, A., Hüther, P., Ashraf, H., Prodanov, T., et al.: Long-read sequencing and structural variant characterization in 1,019 samples from the 1000 genomes project. bioRxiv pp. 2024–04 (2024)

35. Sibbesen, J.A., Eizenga, J.M., Novak, A.M., Sirén, J., Chang, X., Garrison, E., Paten, B.: Haplotype-aware pantranscriptome analyses using spliced pangenome graphs. Nature Methods 20(2), 239–247 (2023)

36. Sirén, J., Monlong, J., Chang, X., Novak, A.M., Eizenga, J.M., Markello, C., Sibbesen, J.A., Hickey, G., Chang, P.C., Carroll, A., et al.: Pangenomics enables genotyping of known structural variants in 5202 diverse genomes. Science 374(6574), abg8871 (2021)

37. Yang, J., Chaisson, M.J.: Tt-mars: structural variants assessment based on haplotype-resolved assemblies. Genome Biology 23(1), 110 (2022)

