## Supplementary Material for "Pangenome graph augmentation from unassembled long reads"

*Luca Denti, Paola Bonizzoni, Brona Brejova, Rayan Chikhi,*
*Thomas Krannich, Tomas Vinar, and Fereydoun Hormozdiari*

### Running times and memory usage

Our approach required 20 minutes and less than 8GB of RAM to augment the $n = 1$ graph (computed with the GNU's `time` utility). When considering the $n = 32$ graph, running time increased to 4.5 hours and the memory consumption to 33GB (peak). The running time of our approach are mostly dominated by the global alignment of each consensus against the (sub)paths of the graph. This indeed required almost 3.2 hours for the $n = 32$ graph. The second most expensive step is the construction of the FMD-index which required 1 hour and 33GB of RAM. Just for comparison, `Minigraph-Cactus` pipeline required 7 hours without considering the assembly step.

### Supplementary Figures and Tables

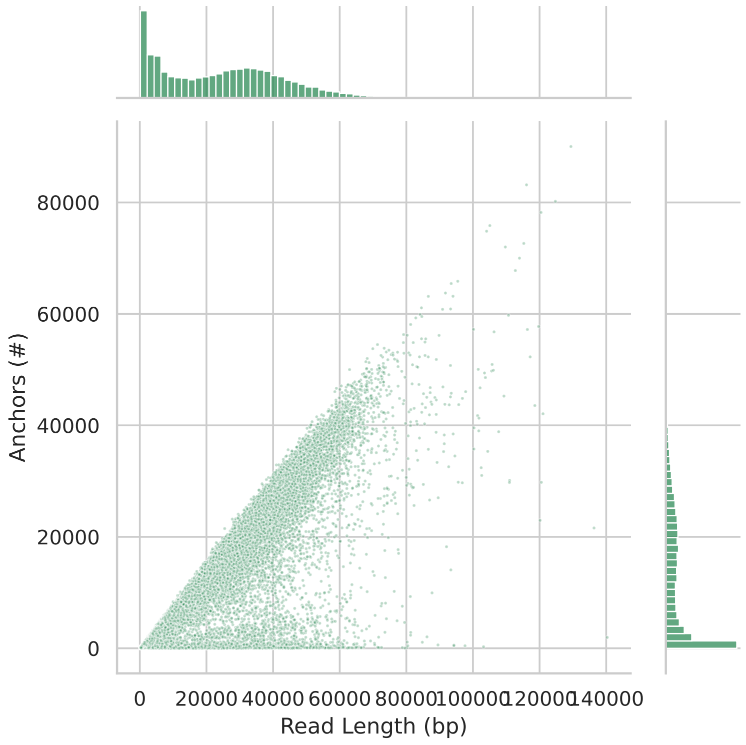


**Fig. A.** Number of anchors and read length distribution in Oxford Nanopore reads (all reads).

| #Samples | Chromosome | Close pairs | Total pairs | Ratio |
|---|---|---|---|---|
| 1 | chr1 | 176650517 | 176705266 | 99.97 |
| 1 | chr11 | 104267678 | 104292462 | 99.98 |
| 1 | chr20 | 153458927 | 153497416 | 99.97 |
| 1 | All | 434377122 | 434495144 | 99.97 |
| 2 | chr1 | 174796985 | 174851673 | 99.97 |
| 2 | chr11 | 103051279 | 103076633 | 99.98 |
| 2 | chr20 | 151434537 | 151473804 | 99.97 |
| 2 | All | 429282801 | 429402110 | 99.97 |
| 4 | chr1 | 172132217 | 172186612 | 99.97 |
| 4 | chr11 | 101337860 | 101364039 | 99.97 |
| 4 | chr20 | 148827057 | 148867766 | 99.97 |
| 4 | All | 422297134 | 422418417 | 99.97 |
| 8 | chr1 | 168085699 | 168141032 | 99.97 |
| 8 | chr11 | 98754037 | 98781590 | 99.97 |
| 8 | chr20 | 145014441 | 145056985 | 99.97 |
| 8 | All | 411854177 | 411979607 | 99.97 |
| 16 | chr1 | 164495172 | 164551620 | 99.97 |
| 16 | chr11 | 96503057 | 96531722 | 99.97 |
| 16 | chr20 | 141572876 | 141616767 | 99.97 |
| 16 | All | 402571105 | 402700109 | 99.97 |
| 32 | chr1 | 159394553 | 159453113 | 99.96 |
| 32 | chr11 | 93385945 | 93416214 | 99.97 |
| 32 | chr20 | 136922211 | 136968520 | 99.97 |
| 32 | All | 389702709 | 389837847 | 99.97 |

**Table A.** Total number of pairs of consecutive anchors on the T2T reference and number of these pairs whose distance is $\leq$ 100bp (`Close pairs` columns). Numbers are broken down by pangenome graph (with increasing number of samples) and by chromosome.

| | #contigs | LargestContig | TotalLen | RefLength | N50 | L50 |
|---|---|---|---|---|---|---|
| **Haplotype 1** | 406 | 1 669 825 | 67 411 189 | 66 210 255 | 337 152 | 52 |
| **Haplotype 2** | 347 | 1 423 908 | 61 386 010 | 66 210 255 | 284 759 | 56 |

**Table B.** Quality assessment for the `chr20` assemblies produced by `hifiasm` from simulated 10x HiFi sample. Assessment has been performed using `quast`.

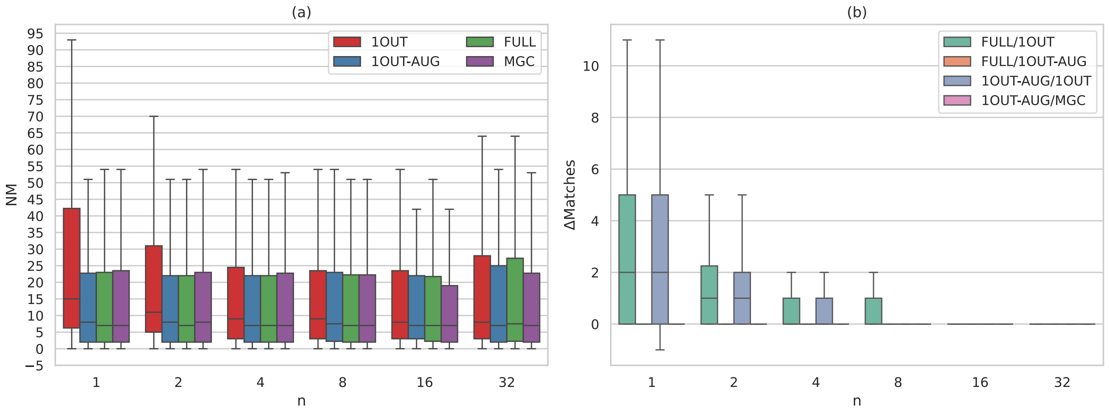


**Fig. B.** Results on alignment accuracy (full alignments). (a) Distribution of the edit distances reported by `minigraph` (`NM` field) when aligning simulated reads to the 4 graphs considered in our analysis (lower is better). (b) Pairwise comparison of the alignments in terms of difference in the number of residue matches ($\Delta Matches$) reported by `minigraph`. In each plot, the results are broken down by the number $n$ of individuals included in the original pangenome graph (x axis). For the sake of clarity, outliers are not plotted. `FULL`: graph built from $n+1$ individuals, `1OUT`: graph built from $n$ individuals, `1OUT-AUG`: `1OUT` augmented with the new individual, `MGC`: assembly-based graph built by `Minigraph-Cactus`).
